# Machine learning-based development of Gadolinium binding peptides

**DOI:** 10.64898/2025.12.14.694215

**Authors:** Nir A. Dayan, Makayla Long, Nicolas Scalzitti, Iliya Miralavy, Daniel Holmes, Mark Kocherovsky, Wolfgang Banzhaf, Assaf A. Gilad

## Abstract

Gadolinium-based contrast agents (GBCAs) are indispensable tools in magnetic resonance imaging (MRI), yet their clinical use is limited by non-specific tissue accumulation, low molecular specificity, and safety concerns. Protein and peptide scaffolds provide a promising alternative because they can bind metal ions with high selectivity and enable precise molecular targeting. However, identifying short peptide motifs with optimal gadolinium (Gd³⁺) coordination and high relaxivity remains a major challenge.

Here, we used a machine-learning-driven peptide evolution platform, the Protein Optimization Engineering Tool (POET), to design and optimize short Gd-binding motifs that enhance longitudinal relaxivity (r₁). Two algorithmic strategies were tested; motif-based and regular-expression-based representations, both were trained on an initial set of 74 twelve-amino-acid peptides derived from natural EF-hand scaffolds. Through two rounds of directed evolution and experimental screening, we identified peptides with up to a 24% increase in r₁ ratio compared with the best natural EF-hand, and a 55% improvement in absolute r₁ after removal of unbound Gd.

Further analysis revealed that peptides exhibiting higher relaxivity generally possessed a more negative net charge and lower isoelectric point than the buffer pH, indicating stronger electrostatic stabilization of Gd³⁺. Sequence enrichment analysis showed that acidic and small polar residues, particularly aspartic acid, glycine, and threonine, were selectively favored during evolution, while bulky hydrophobic and basic residues were depleted. These compositional trends align with improved solubility and enhanced metal coordination.

Together, these results demonstrate a generalizable framework that integrates computational evolution with biophysical screening to discover new biologically derived Gd-binding motifs. This approach provides a scalable route to engineer responsive, tunable, and biocompatible MRI contrast tags for precision imaging and molecular diagnostics.

## INTRODUCTION

Gd is a rare earth element (REE) and one of fifteen lanthanide group elements. Gd’s electron configuration makes it highly reactive, and it enhances proton relaxation to create a high magnetic moment making it an ideal element for an MRI contrast agent^1,2^. Since 1988, gadolinium-based contrast agents (GBCAs) have been developed to improve stability and reduce toxicity, but they have remained the predominant choice as high-quality contrast imaging agents^3^. The negative aspect of GBCAs is that they are a broad contrast agent administered into the blood stream in order to disperse throughout the entire body, which has been shown to cause other adverse effects such as accumulation in skin, muscle, bone, and brain^4–8^. One of the reasons Gd accumulates in the body is that the Gd cation (Gd^+3^), when unchelated, has biophysical characteristics similar to Ca^+2^, and several studies have investigated the potential mimicry effects this may have on body systems. Research found that Gd^+3^ ions increased mitochondrial fluidity, resulting in promotion of apoptosis in mice livers, contributing to further suspicions of unknown toxicities of Gd in the body^9^. Previous research has also shown that the binding of GdCl_3_ to tubulin of NIH-3T3 cells might inhibit the assembly of tubulin or depolymerize microtubules in cells, while the GBCA gadolinium diamide (gadodiamide) showed much lower levels of toxicity^10^. That being said, studies have shown GBCAs are not as inert to protein binding as many assume, demonstrating that more work must be done to understand the implications that weak but broad protein binding may have in vivo^11^. Therefore, it is extremely desirable to develop GBCAs that are more efficient, reducing the amount of potential harmful metal administered to patients.

Biologics are substances and products derived from living organisms that are used to prevent, diagnose, or treat a variety of health conditions^12^. In randomized control trials, biologics have demonstrated an ability to improve quality of life for patients suffering from joint damage, skin lesions, and gastrointestinal inflammation^13,14^. Not only have biologics demonstrated potential to be effective pharmacological agents in chronic disease, but they also present opportunities for increased precision and tailoring to patient needs. It is highly desirable to track biologics in vivo as part of theranostics therapies, since it is important to know whether the biologic reached its target in the body. GBCAs have been used to track biologics using MRI^15^, with most efforts targeting chemical conjugation of GBCA to the biologic. However, attaching Gd chelates to antibodies, for example, often produces heterogeneous labeling, leading to variable relaxivity, impaired binding, and reduced immunoreactivity. Because each chelate contributes only modestly to the MRI signal, multiple Gd units are required, complicating synthesis, purification, and reproducibility^16,17^.

By contrast, biologics-based scaffolds provide a more controlled alternative. For example, lanmodulin^18^, a recently discovered bacterial protein with picomolar affinity and 10⁸-fold selectivity for lanthanides over Ca²⁺, coordinates lanthanides using EF-hand motifs, short helix–loop–helix structures that typically bind Ca²⁺ in the calmodulin protein^19^. In lanmodulin, four EF hands have been adapted for lanthanide binding, providing a natural, high-affinity scaffold for engineering next-generation GBCAs. Engineered variants such as the single-point mutated LanND-Gd^20^, achieve longitudinal relaxivity of 13.15 mM⁻¹·s⁻¹ at 3 T, significantly higher than that of the clinically used agent Magnevist (4.62 mM⁻¹·s⁻¹ at 3 T).

Another prominent biologics-based platform is the Protein Contrast Agent (ProCA) series^21^, ProCA32^22^, a third-generation protein GBCA based on calmodulin with engineered EF-hand motifs, which achieves r₁ values of ∼33 mM⁻¹·s⁻¹ at 1.4 T and 37 ^0^C, nearly tenfold higher than Magnevist (3.3 mM⁻¹·s⁻¹ under identical conditions). ProCA32 can also be fused with targeting moieties (e.g., HER2, EGFR, VEGFR) without compromising metal coordination, enabling high-sensitivity molecular imaging of cancers, fibrosis, and metastases in vivo.

Together, LanND-Gd and ProCA32 show how protein scaffolds can deliver an order-of-magnitude boost in relaxivity while preserving targeting and stability. By tightly coordinating Gd³⁺ within EF-hand motifs, these biologics minimize metal release and enable fusion with recognition domains, making them potentially powerful platforms for high-sensitivity, clinically relevant molecular imaging.

Despite these advances, current protein-based GBCA remain limited by the need to optimize metal-binding motifs for maximal relaxivity in a systematic, predictive manner. Most efforts rely on single-point mutations or rational design, which can be laborious and may miss nonintuitive sequence combinations that could further enhance performance. There is a clear need for computational tools that can efficiently explore the vast sequence space of short motifs, learn sequence–function relationships, and propose variants with improved relaxivity and binding properties.

Here we address this gap by applying POET^23^ (Protein Optimization Engineering Tool), a genetic programming–based algorithm, to systematically explore and optimize lanthanide binding peptides. POET has already been successfully applied to optimize short amino acid motifs for chemical exchange saturation transfer (CEST) MRI applications^24–26^, demonstrating its ability to efficiently explore sequence space and uncover nonintuitive variants with enhanced performance. POET has two versions which use different methods to recognize amino acid sequences. The original POET (called here POET_motif_^23^) in which individuals are composed of amino acid subsequences paired with corresponding weights, and an extension of POET (called here POET_regex_^25^) where individuals are represented by a list of regular expressions (regex) and corresponding weights.

Our objectives in the present work were to (i) discover short motifs with improved r₁ relative to natural EF-hand using a high-throughput assay, and (ii) compare POET_motif_ and POET_regex_ to assess efficiency and diversity of outputs. More broadly, we aimed to establish a pipeline for motif discovery that could inform the design of next-generation protein-based MRI contrast agents.

## METHODS

### Peptide Synthesis and Preparation

The peptides were purchased from Genscript USA Inc. (Piscataway, NJ). HEPES ((4-(2-hydroxyethyl)-1-piperazineethanesulfonic acid)) and GdCl_3_ were purchased from Thermo Scientific Chemicals, Waltham, MA USA. HEPES was chosen because other buffers containing phosphates tend to form Gd-phosphate sediments.

### MRI measurements

The ability of a contrast agent (CA) to produce MRI contrast is quantified by relaxivity, which is the CA’s ability to change the relaxation rate of a solution as a function of its concentration. To measure relaxivity one must measure the longitudinal relaxation time (T_1_) of several dilutions of the CA, and extract the slope from linear regression of the following equation

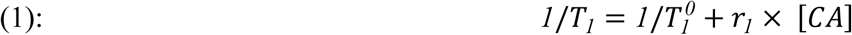

Where *T*_1_ and 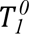 are the longitudinal relaxation time of the solution with and without the CA respectively. r_1_ is the longitudinal relaxivity and [CA] is the GdCl_3_ concentration of the CA measured by inductively coupled plasma (ICP) (Varian 710-ES). ICP samples were prepared by nitric acid digestion.

Peptides were dissolved in HEPES 50mM and GdCl_3_ 0.2mM pH 5.3 and incubated r_1_ = r_1_ (peptide)/ r_1_ (GdCl_3_)) measurements the samples were diluted to measure their T_1_ values, along with GdCl_3_ that did not have any peptide. To perform more high-throughput measurements, Gd concentrations were not measured for these peptides. Instead, the peptides’ r_1_ were divided by the peptide-free Gd r_1_. For absolute relaxivity, MRI was conducted 42 hours after incubation with GdCl_3_ and two dialysis cycles.

To create a series of dilutions, an epMotion 5073 liquid handler was used to ensure pipetting accuracy. The sample, along with 6 or 7 sample dilutions and a blank solution, were pipetted into a 396 plate cut to fit inside the MRI magnet bore, sample volume inside the wells was ∼100 microliters. 7 columns along the phase axis and 5 rows along the read axis could be measured with a relatively homogeneous field. Two custom fitted plates were stacked on top of each other, enabling measurements of two slices simultaneously.

MRI measurements were performed on a horizontal 30 cm bore 7 T Bruker preclinical MRI with ParaVision 360 v3.2 on a Linux computer running CentOS7, using a B-GA12SHP high performance gradient insert with built-in shims. A variable repetition time T_1_ map was acquired using a RARE sequence with 9 TRs (33 122 297 497 727 1577 2777 4809 12496) ms/TE 6.88 ms, RARE factor = 1, FOV ∼ 40x25 mm, with the matrix adjusted to ensure a spatial resolution of 0.25x0.25 mm, slice thickness = 1.2 mm, 1 or 2 slices, echo spacing was 6.88 ms. This is correlated to measurement times of 40 to 60 minutes. Finally, the T_1_ for each well was measured using the image sequence analysis module in Paravision, then r_1_ ratio or absolute r_1_ relaxivity were calculated.^27^

### Peptide evolution using machine learning

Two versions of the POET algorithm^28^ were compared for their ability to create peptides that can shorten the T_1_ relaxation of water and have higher r_1_ ratio. POET_motif_ and POET_regex_., the former using motifs represented as substrings, and the latter using regular expressions (regex), a formal way of defining a set of character sequences^29^. A motif/regex was classified as beneficial when it had r_1_ ratio > 1, which corresponds to the r_1_ ratio of GdCl_3_ without a peptide. POET uses a form of supervised training called Genetic Programming. Each POET trial generates a set of rules that determine r_1_ ratio by checking the composition of a proposed peptide sequence. If a given motif (or regex) is found, the corresponding weight is added to a sum value, in this case r_1_ ratio. The rulesets interact through recombination and mutation over timesteps (called *generations*), and once all generations are completed, the most accurate model is returned for peptide generation. Both POET variants were trained with 74 12-amino acid long peptides incubated with GdCl_3_.

The r_1_ of the peptides was normalized to the r_1_ of the buffer with the same amount of GdCl_3_.

### Nuclear magnetic resonance (NMR) measurements

Solution NMR experiments were carried out on a 600 MHz Bruker Avance NEO NMR spectrometer equipped with a 5 mm Prodigy TCI cryoprobe. ^1^H NMR experiments were run at 25°C using the vendor supplied water suppression pulse sequence (noesygppr1d) to measure ^1^H spectra of 1mM peptide of EF3 and CaBM^30^. Both peptides were suspended in HEPES 50mM with 10% D_2_O for locking, shimming, and chemical shift referencing with or without 5μM GdCl_3_.

## RESULTS AND DISCUSSION

### Peptide evolution

Both POET algorithms were applied to evolve Gd-binding peptides with enhanced MRI relaxivity. Figure 1 and S1 show plots of the r_1_ ratio of the peptides predicted by POET_motif_ and POET_regex_ over two POET cycles dubbed Epoch 1 and 2. In Figure 1 the red dashed line represents the r_1_ ratio of EF hand 3 motif (EF3) (r_1_=1.86), a 12-amino-acid from Lanmodulin that showed the highest r_1_ ratio from the training data. For the POET_motif_ in Epoch 1, only three peptides showed better contrast than the EF3 peptide; however, in Epoch 2, 9 of 10 peptides surpassed the r_1_ ratio of the EF3 hand peptide. For POET_regex_ 7 out of 10 peptides surpassed the r_1_ ratio of EF3 in both Epochs 1 and 2. The best motif found had an r_1_ ratio of 2.3, a 24.3% improvement over the EF3. Peptides displaying an r_1_ relaxivity ratio lower than that of EF3 suggest that POET did not have sufficient data to fully explore the sequence space. Nevertheless, the algorithm successfully identified novel peptides with enhanced relaxivity, demonstrating its capacity to discover functionally improved motifs beyond those found in nature. Next, the correlation between POET’s predictions and MRI measurements was examined. No correlations were observed, indicating that more epochs could improve the correlation.

**Figure 1.**
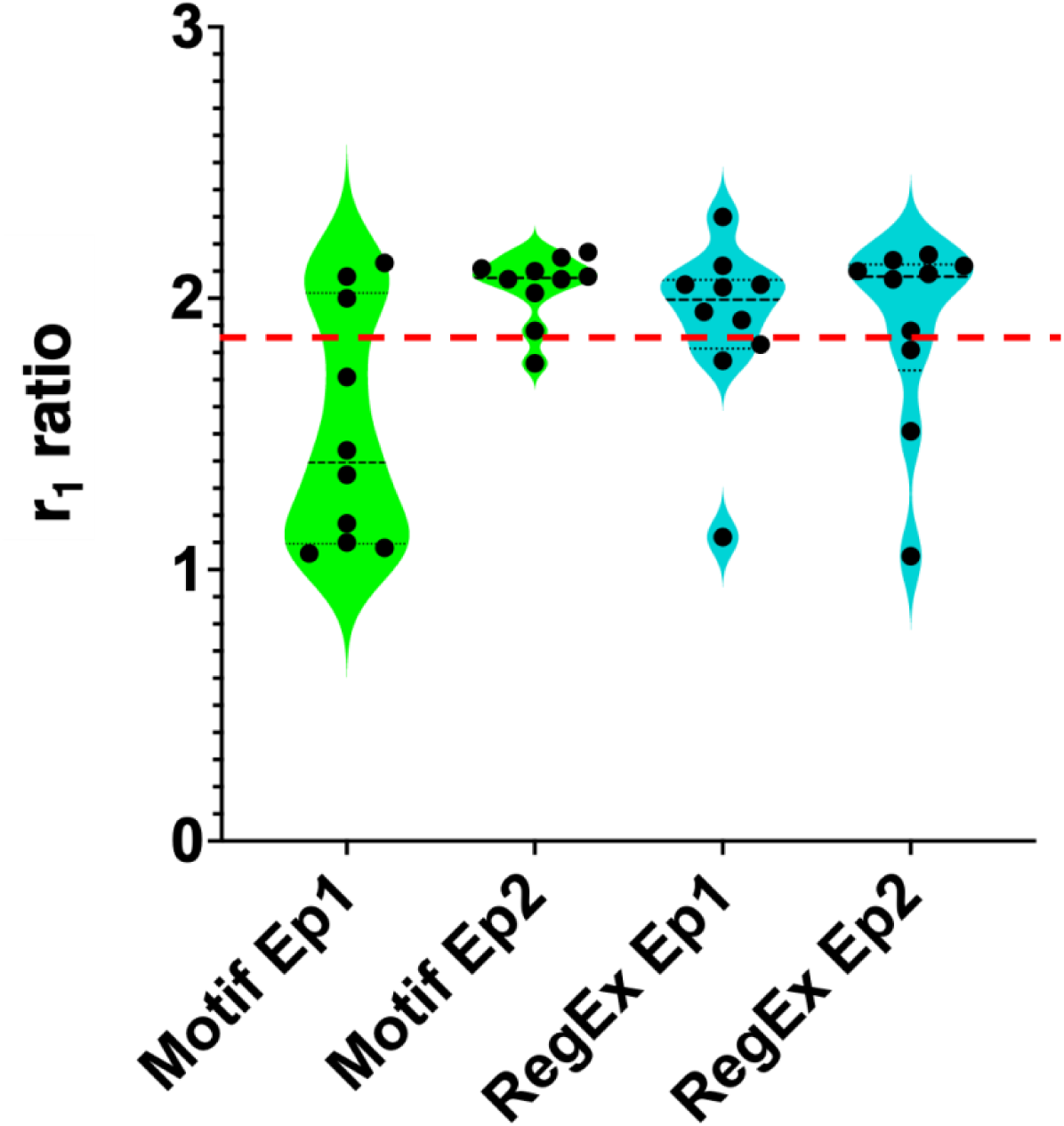
r_1_ ratio of peptides predicted by POET. The red dashed line represents the r_1_ ratio of the EF 3 hand motif (EF3).

### Validation of the Gd-relaxivity

The *r₁ ratio* is a convenient high-throughput metric reflecting the ability of Gd-binding peptides to enhance longitudinal relaxivity; however, it represents combined contributions from peptide-bound and free Gd³⁺ ions. To verify this and to characterize the metal species present prior to dialysis, we performed electron spin resonance (ESR) spectroscopy (Supporting Information, Figure S1). The dialyzed EF3 sample exhibited a broad, downfield-shifted, low-intensity ESR signal characteristic of peptide-bound Gd³⁺. In contrast, the undialyzed EF3 + Gd³⁺ mixture, after subtraction of the free-Gd spectrum, displayed a strong residual central peak and a downfield shoulder. The central peak reflects an incomplete cancellation of the intense free-Gd transition, while the shoulder represents a heterogeneous population of Gd³⁺ interacting with the peptide. The distinct downfield positions of the two traces indicate that the bound Gd³⁺ species present before and after dialysis occupy slightly different electronic environments due to differences in exchange, hydration, and coordination equilibrium. These ESR results show that undialyzed samples contain a large fraction of free Gd³⁺ with a different electronic environment than the peptide-bound species observed after dialysis. Therefore, we measured absolute r₁ after dialysis to confirm that it correlates with the r₁ ratio used in the initial screen.

With this rationale, the absolute relaxivities (r_1_) of Gd-binding peptides were measured after two cycles of dialysis over 48 hours against HEPES buffer to remove excess Gd³⁺, which typically reduced the total Gd concentration from 0.2 mM to ∼20 µM. Peptides showing a range of r_1_ ratios (Table 1) were selected, including EF-hand motifs, a calcium-binding motif (CaBM) peptide, a lanthanide-binding peptide, and peptides predicted by POET. A strong correlation was found between the r₁ ratio and absolute r_1_ (Figure 2; R² = 0.6137, p = 0.0073). These findings indicate that r_1_ ratio is a reliable predictor of absolute relaxivity and can be used as an efficient screening metric to minimize time and material use. Moreover, when comparing absolute r_1_ values, a 42.6% improvement was observed for the best peptide identified in the initial screening (IDCRGTEDDDPN) relative to EF3, versus the 24.3% improvement indicated by its r₁ ratio. The peptide showing the highest absolute r₁ (TQDSDDGMEDED) displayed a 55.2% enhancement over EF3. Taken together, these results demonstrate that POET can successfully identify novel peptides that coordinate Gd³⁺ and achieve relaxivities exceeding those of natural sequences.

**Figure 2.**
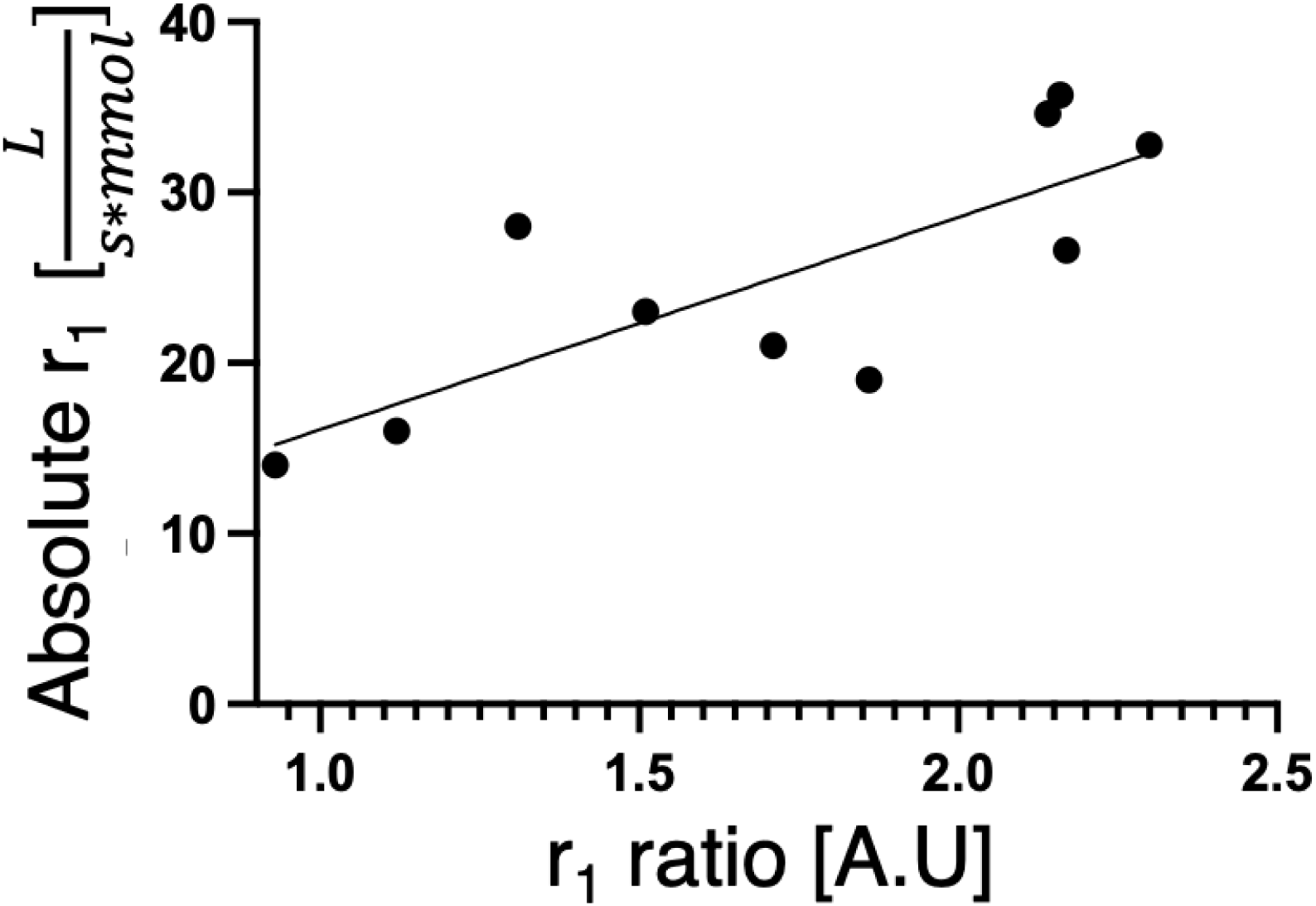
Correlation between the r_1_ ratio and the absolute r_1_. Goodness of fit (R^2^= 0.6137, p-value =0.0073).

**Table 1:** Comparison of absolute r_1_ to r_1_ ratio.

| Source | Peptide sequence | $r_1$ ratio | Absolute $r_1 \left[ \frac{L}{s \cdot mmol} \right]$ |
| --- | --- | --- | --- |
| **CaBM<sup>30</sup>** | DKDGNGYISAAE | 0.93 | 14 |
| **GdCl<sub>3</sub> 0.2mM** |  | 1.00 | 35 |
| **POET regex1** | NSAMADQADDDP | 1.12 | 16 |
| **Lanthanide binding<sup>31</sup>** | DKDGDTIDEREI | 1.31 | 28 |
| **EF 1 hand motif<sup>32</sup>** | DPDKDGTIDLKE | 1.51 | 23 |
| **POET Motif1** | DMMHDKTDCNDT | 1.71 | 21 |
| **EF3<sup>32</sup>** | DPDNDGTLDKKE | 1.86 | 19 |
| **POET regex 2** | SDDNHSDLGDDL | 2.14 | 34.6 |
| **POET regex 2** | TQDSDDGMEDED | 2.16 | 35.7 |
| **POET motif 2** | GDDEDDQCHQDG | 2.17 | 26.6 |
| **POET regex 1** | IDCRGTEDDDPN | 2.30 | 32.8 |

### Analysis of peptide’s isoelectric point and net charge

After two Epochs, an extensive analysis was performed to evaluate POET’s predicted peptides. First, the net charge and the isoelectric point (IP) at pH 5.3 were calculated for each peptide and plotted against their r_1_ ratio. The results in Figure 3 show that most peptides with an r_1_ ratio better than free-Gd had a negative net charge, and usually the r_1_ ratio improved with more negative net charge, since Gd has a 3+ positive charge this makes sense. For the IP, peptides with values under 5.3 had higher r_1_ ratios, since the buffer’s pH was 5.3, only peptides with lower IP had negative charges that could stabilize the Gd^3+^ ions. The results suggest that for further evolution, the peptides’ net charge should be negative, and their IP should be lower than the buffer’s pH, both conditions easily screened using POET. It is possible that some motifs and regular expressions were neglected by POET because the net charge of the molecule was too high, so it could be beneficial for future Epochs to use only the negatively charged peptides. It could also be useful to choose the buffer pH to be as high as possible, leading peptides that have higher IP to have higher r_1_ values, making the dataset more helpful for learning in POET.

**Figure 3.**
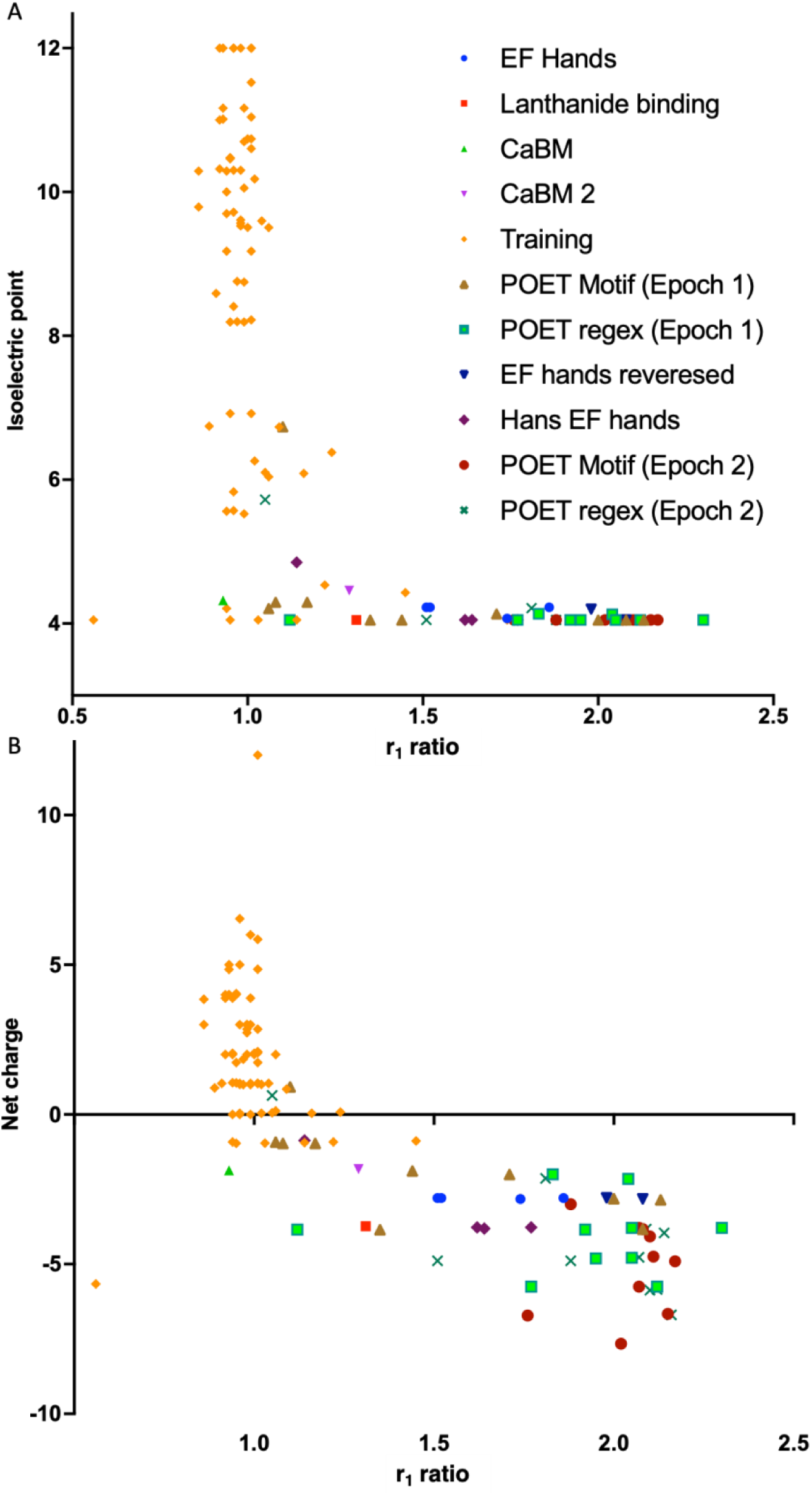
A) Calculated net charge of a peptide plotted against its r_1_ ratio. B) Calculated isoelectric point of a peptide plotted against its r_1_ ratio. The isoelectric point was calculated for pH 5.3. The different groups are listed in Table S1.

### Paramagnetic relaxation enhancement (PRE)

PRE measurements were performed to validate that Gd binds to the peptides, and to investigate which amino acids are more involved in the binding. A comparison was made between similar EF-hands, EF3^32^, known to bind lanthanides, and CaBM^30^, known to bind Calcium.

Since Gd^3+^ is a paramagnetic ion that significantly shortens both the T_1_ and T_2_ ^33^ of any NMR active nuclei within approximately 25 angstroms of the binding site, NMR was used to measure ^1^H spectra of 1mM peptide of EF3 and CaBM with or without 5μM GdCl_3_. As shown in Figure 4, EF3 displays a more significant increase in the linewidths of all proton peaks with a concomitant reduction in intensity due to paramagnetic relaxation than those of CaBM. PRE will be more pronounced for those protons proximal to the binding site than for those distal, which is why EF3 displays more significant broadening of the NH peaks of Asp-D, Gly-G, Leu-L, and Thr-T (7.9-8.4 ppm) than other proton resonances in the peptide. CaBM also displays broadening for Asp-D, Gly-G, Ser-S, and Glu-E, but to a lesser extent, and there is comparatively little relaxation enhancement for the aliphatic protons of leucine (0.95 and 1.6 ppm) thereby indicating that these protons are more distal to the Gd binding sites. The number of NH resonances broadened into the baseline is higher for EF3 than CaBM which can be attributed to a higher percentage of amino acids binding to Gd for EF3 than CaBM.

**Figure 4.**
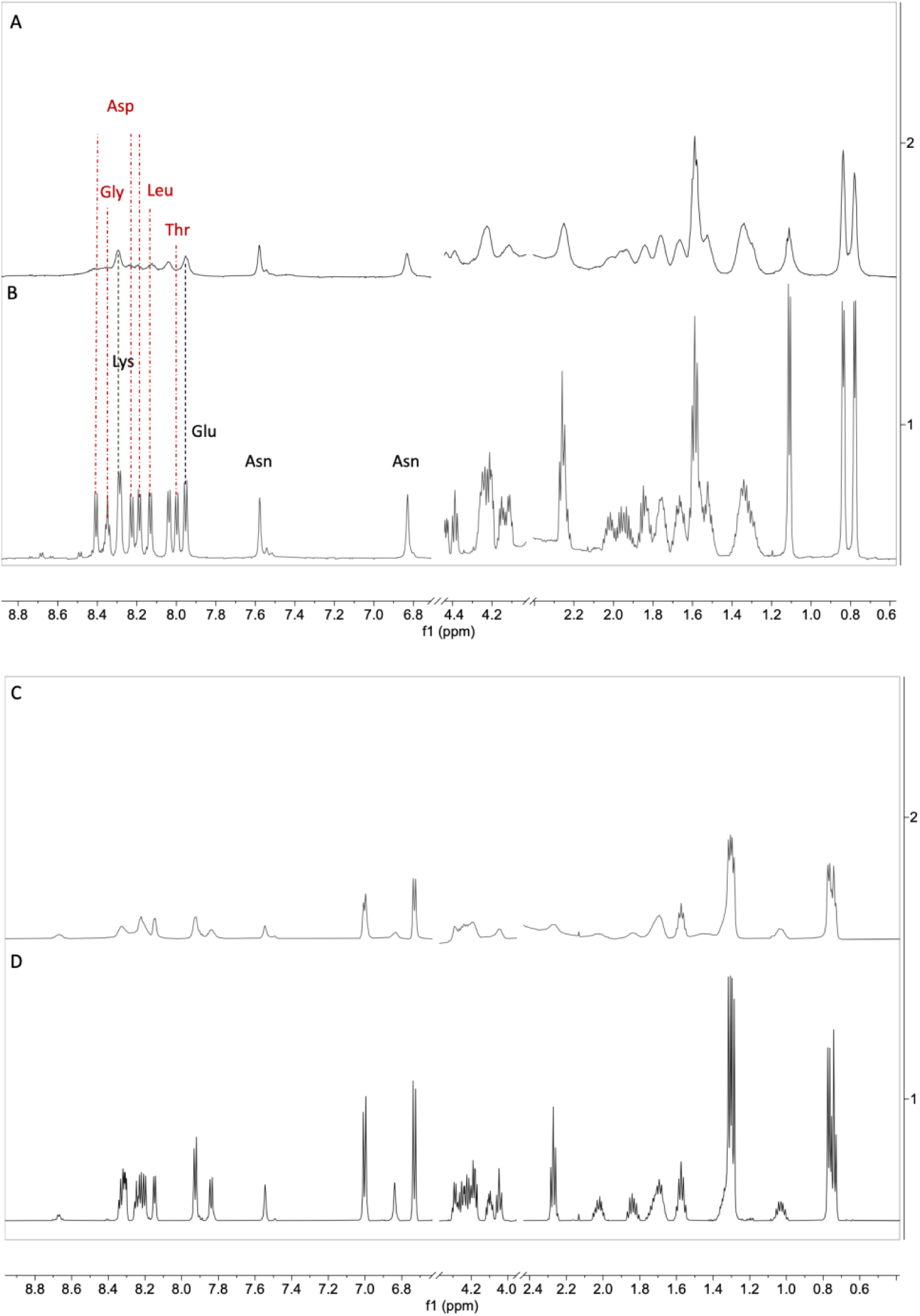
^1^H NMR spectra of (A) EF3 DPDNDGTLDKKE (1mM peptide) with 5μM GdCl_3_. (B) EF3 (1mM peptide). The vertical lines connect between the peaks to show the line broadening when there’s presence of GdCl_3_. The red dot-dashed line connects the peaks that were broadened to a higher extent, while the black dashed line connects the peaks that were less broadened. (C) The Ca binding motif (CaBM) DKDGNGYISAAE (1mM peptide) with 5μM GdCl_3_ and (D) CaBM (1mM peptide).

### Amino acid distribution

The performances of POET_motif_ and POET_regex_ were evaluated and compared by the amino acid composition of each algorithm and Epoch. Figure 5A shows the distribution of the different amino-acids classes within the resulting peptide sequences, Acidic (D, E), Basic (K, H, R), Aliphatic (A, G, I, L, P, V), Aromatic (F, W, Y), Hydroxylic (S, T), Sulfur-containing (C, M) and Amidic (N, Q). Figure 5B illustrates the individual amino acid distribution. Importantly, the initial training dataset exhibited a nearly uniform distribution of amino acid residues, providing an unbiased starting point for the evolution performed by POET.

**Figure 5.**
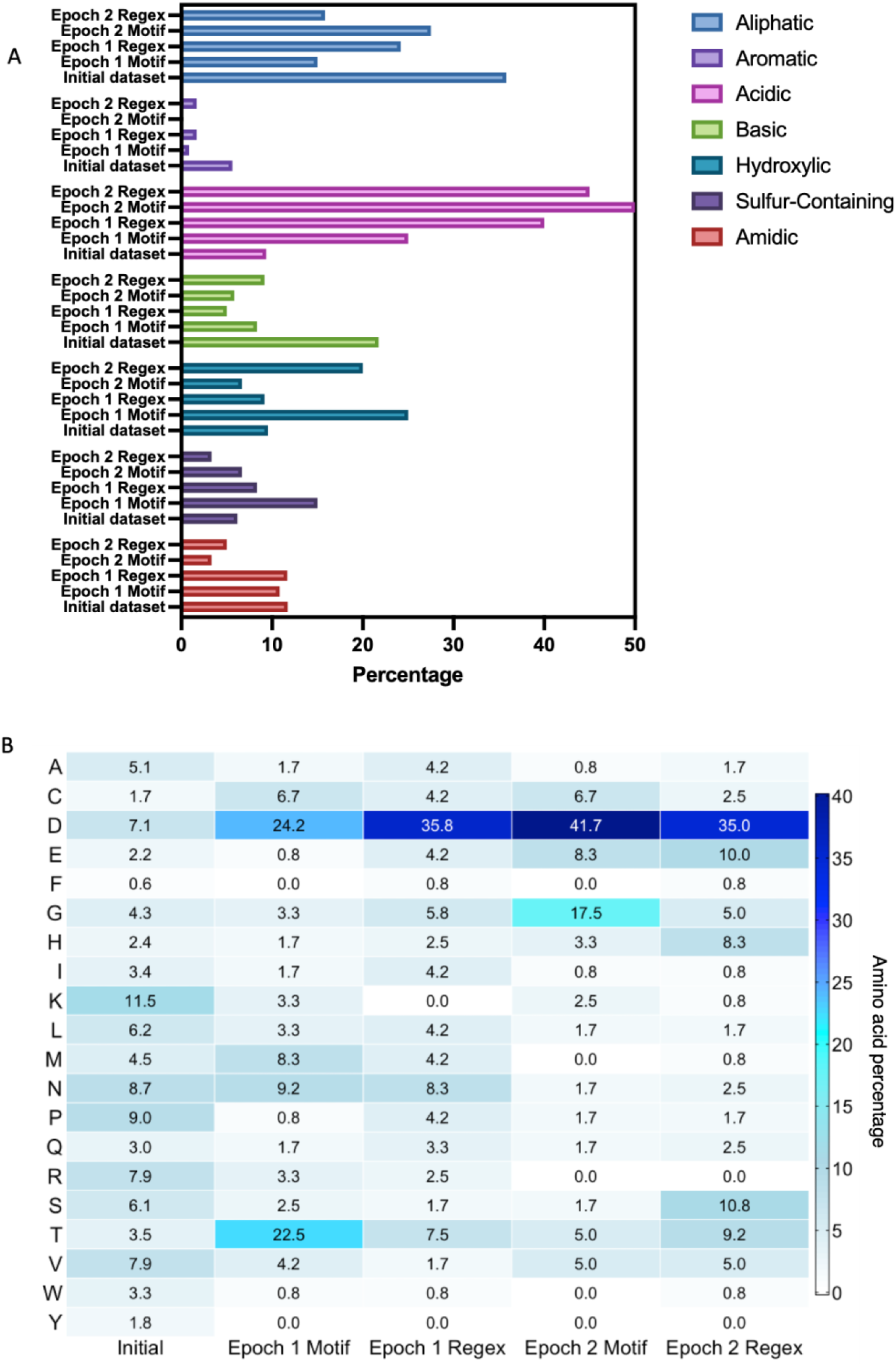
The amino acid distribution in the training data, Epoch 1 and 2, with either motif or regular expressions (Regex) POET. A. Column bar representation of the amino acid categories distribution across the datasets. B. Heatmap of the amino acid’s percentage in the different datasets.

Across all groups, there was a consistent and pronounced enrichment of acidic residues (particularly aspartic acid, D) and a marked depletion of basic amino acids (notably lysine, K, and arginine, R). This suggests a strong selection for negatively charged residues capable of coordinating with the positively charged Gd³⁺ ions, and simultaneous avoidance of positively charged residues to minimize electrostatic repulsion. Aspartic acid emerged as the dominant residue in both, motif and regex patterns during both evolutionary epochs, comprising over 24% of amino acids in all of POET’s outputs. This selection mirrors the composition of many natural EF-hand motifs, which often contain multiple aspartic acids involved in metal ion coordination. While there was a general reduction in non-polar aliphatic residues, one exception was glycine (G), which increased significantly in Epoch 2 motifs. Glycine’s small side chain likely contributes to increased peptide flexibility and solubility, properties that may be advantageous in a hydrophilic or dynamic binding context. The hydroxylic class (serine and threonine) showed moderate enrichment, particularly threonine (T), which rose sharply in Epoch 1 Motif. These residues support hydrogen bonding and surface polarity, reinforcing the hydrophilic nature of the evolving peptides.

The aromatic and basic classes, especially phenylalanine (F), tryptophan (W), and lysine (K), were consistently depleted across all outputs, indicating a preference against bulky, hydrophobic, or positively charged side chains. Sulfur-containing and amidic residues (C, M, N, Q) exhibited more subtle variations, with no strong trend in either direction.

Consistent with these compositional trends, PRE measurements confirmed that several of the amino acids enriched by POET are indeed directly involved in Gd³⁺ coordination. In particular, residues such as aspartic acid (D), glycine (G), and threonine (T), which showed strong enrichment in the evolved sequences, were among those that exhibited significant signal broadening in EF-hand peptides upon Gd binding.

Notably, the inclusion of all 20 amino acids in the training set allowed POET to explore a broad sequence space during motif and regex discovery, enabling the algorithm to converge on a highly specific and chemically coherent set of residue preferences aligned with metal-binding functionality.

## CONCLUSION

We applied POET to expand and optimize short lanthanide-binding peptide motifs and demonstrate a practical, data-driven route to improve Gd-mediated T_1_ contrast. Starting from an experimental library of 74 sequences and using iterative POET cycles, we identified novel peptides with substantially enhanced relaxivity, the best POET-predicted motif yielded an r₁-ratio of 2.30 (∼24% higher than the best initial EF-hand, EF3), and top peptides measured after dialysis showed up to ∼55% higher absolute r₁ than EF3. The screening metric (r_1_ ratio) correlated with absolute relaxivity (R² ∼ 0.61, p ∼ 0.007), validating the high-throughput screening approach, and PRE spectra support that Gd³⁺ is in proximity to peptide residues. Sequence analysis revealed strong enrichment of acidic residues (notably Asp), consistent with electrostatic and coordinating roles in Gd binding.

These findings show that coupling POET with a high-throughput r₁ screening can efficiently explore short-motif sequence space and produce superior candidates; and that running complementary POET recognition modes (motif and regex) yields both rapid consensus improvements and broader, nonintuitive solutions. Some discovered peptides could be explored as standalone contrast agents; future work will evaluate motif transferability into full proteins (e.g., Lanmodulin) and biologics (e.g., antibodies). Overall, POET combined with a high-throughput pipeline provides the foundation for designing peptide tags and for guiding the development of next-generation biologics-based Gd MRI contrast agents with improved sensitivity and translational potential.

## DATA ACCESSIBILLITY STATEMENT

POET’s code can be accessed freely at https://gitlab.com/NicolasScalzitti/poet_regex and https://github.com/elemenohpi/POET

## ACKNOWLEDGEMENTS

We would like to thank ICER at MSU for giving us access to the HPCC system for all our computational work. We would also like to thank - Dr. Christiane Mallett from the advanced molecular imaging facility at MSU for her assistance with MRI setup and analysis; and Prof. John L. McCracken for his assistance with ESR measurements and analysis. We thank Dr. Aidan Murphy for commenting on an earlier version of the manuscript. A.A.G acknowledges financial support from the NIH/NIBIB: R01-EB031008; R01-EB031936. A.A.G. and W.B. acknowledge support from R01-EB030565.

## Abbreviations

REE: Rare earth elements
GBCA: Gadolinium-Based Contrast Agents
ProCA: Protein Contrast Agent
POET: Protein Optimization Engineering Tool
EF3: EF hand 3 motif
CaBM: Calcium Binding Motif
RegEx: Regular Expressions
IP: Isoelectric Point
PRE: Paramagnetic Relaxation Enhancement

