## Supplementary Material for "Machine learning-based development of Gadolinium binding peptides"

Electron spin resonance (ESR) spectroscopy[1] was used to assess the relative contributions of bound and free $Gd^{3+}$ ions in solution. EF3 peptide (740 µM) was incubated with $GdCl_3$ under two conditions: (i) 1 mM $GdCl_3$ followed by dialysis to remove unbound metal and (ii) 250 µM $GdCl_3$ without dialysis. Measurements were performed in 50 mM HEPES at 130 K on an X-band continuous-wave ESR spectrometer. Figure S1 shows the comparison of these spectra. The black trace represents EF3 incubated with $Gd^{3+}$ and subsequently dialyzed to remove free ions, while the blue trace corresponds to the spectrum subtraction of (undialyzed EF3 incubated with Gd) minus (free $Gd^{3+}$ spectrum of the same concentration). Thus, the figure does not display the raw undialyzed EF3 or free $Gd^{3+}$ spectra individually, but rather the difference spectrum isolating the contribution of $Gd^{3+}$ bound to the peptide. The dialyzed sample (black trace) exhibits a broad, downfield shifted low-intensity feature characteristic of peptide-bound $Gd^{3+}$, whose restricted tumbling and altered electronic environment broaden and shift the signal. A small central peak remains, reflecting a minor amount of free $Gd^{3+}$ that persists after dialysis.

The subtracted spectrum (blue trace) retains both a strong central peak and a downfield shoulder. The intense central peak results from incomplete cancellation of the free-Gd transition, which can differ in amplitude or linewidth between samples and therefore does not fully subtract. The downfield shoulder arises from peptide-bound or partially bound $Gd^{3+}$ in the undialyzed mixture. This bound population is not identical to the dialyzed species; it is more heterogeneous, influenced by higher free-Gd concentration and dynamic exchange with bulk solvent, leading to a slightly different ESR position and shape. Together, these data confirm that dialysis removes most unchelated $Gd^{3+}$ and isolates a stable peptide-bound population, validating the conditions used for subsequent relaxivity measurements.

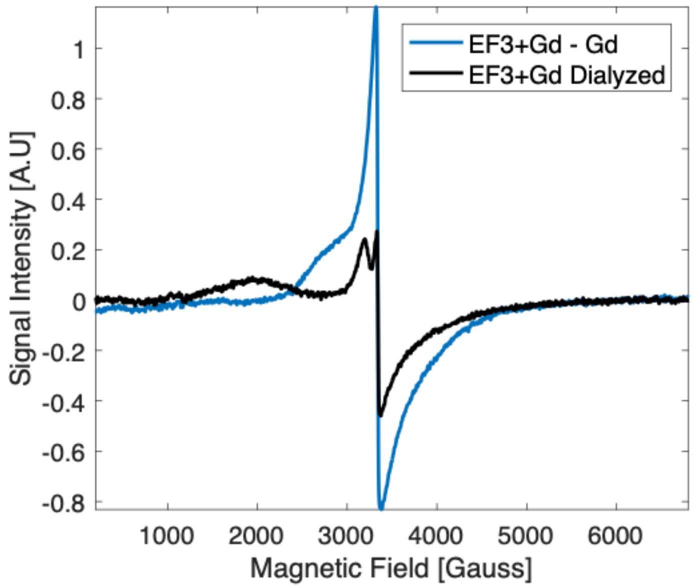


**Figure S1.** Electron spin resonance (ESR) spectra of EF 3 hand motif (EF3) peptide incubated with $Gd^{3+}$ under two conditions. The black trace represents EF3 (740 μM) incubated with 1 mM $GdCl_3$ and dialyzed to remove unbound metal. The blue trace shows the spectrum subtraction of EF3 (740 μM) incubated with 250 μM $GdCl_3$ without dialysis and free $Gd^{3+}$ of the same concentration. Measurements were performed in 50 mM HEPES at 130 K using X-band continuous-wave ESR. The dialyzed sample exhibits lower intensity and broader lines, consistent with peptide-bound $Gd^{3+}$ and removal of unchelated ions.

Peptide dataset and results

Table S1 contains the entire dataset, peptides 1 to 74 were used as the initial training for POET predictions, dubbed Epoch 1. Epoch 2's training was comprised of the initial training set, Epoch 1, the reversed sequence of EF hands 3 and 4, and additional EF hands (peptides 97-100 in Table S1). The reversed sequences of 2 EF hand motifs were measured to check if POET can consider the entire dataset's reverse sequence as another dataset, this way doubling the amount of available data. However, MRI results showed that the regular and reversed sequences had different $r_1$ ratios. Figure S2 provides a graphic representation of the results.

***Table S1. Complete list of peptides***

| | Sample | Sequence | $r_1$ ratio |
|---|---|---|---|
| **1** | EF hand 1[2] | DPDKDGTIDLKE | 1.51 |
| **2** | EF hand 2[2] | DPDKDGTLDAKE | 1.52 |
| **3** | EF hand 3[2] | DPDNDGTLDKKE | 1.86 |
| **4** | EF hand 4[2] | NPDNDGTIDARE | 1.74 |
| **5** | Lanthanide binding[3] | DKDGDTIDEREI | 1.31 |
| **6** | CaBM 2[4] | DKDGDGTITTKE | 1.29 |
| **7** | CaBM[5] | DKDGNGYISAAE | 0.93 |
| **8** | Training | DAPSFAKVVPNR | 0.39 |
| **9** | Training | KVNFNKAVSNLK | 0.40 |
| **10** | Training | LVKPMNKRWVRM | 0.40 |
| **11** | Training | LSNRRGREQYAG | 1.01 |
| **12** | Training | GIFKTTKCKHNS | 0.86 |
| **13** | Training | KVIRYVVAPMKL | 0.86 |
| **14** | Training | CHLKDLRKMGLR | 0.99 |
| **15** | Training | VKKYVTPGAPVY | 0.98 |
| **16** | Training | QPRMWVNVYTCY | 0.97 |
| **17** | Training | NQHTPPKPLNNA | 0.97 |

| | | | |
|---|---|---|---|
| **18** | Training | LHKDRRKPLPLP | 0.92 |
| **19** | Training | QLGVHRNPRWNQ | 1.01 |
| **20** | Training | HLRLNKNRMQKS | 1.01 |
| **21** | Training | NSSNHSNNMPCQ | 1.09 |
| **22** | Training | HQWHRHRPPIRR | 0.96 |
| **23** | Training | RWHHVNVLWQDR | 0.98 |
| **24** | Training | DNPHQLLPVNWR | 0.89 |
| **25** | Training | WFGLQRHLKKKD | 0.94 |
| **26** | Training | GQRWLYKMKDSM | 0.94 |
| **27** | Training | LDHTWGKWGHQS | 0.95 |
| **28** | Training | PVARKVVQICHP | 0.98 |
| **29** | Training | DPMRVRPWYVAR | 1.00 |
| **30** | Training | PGGVPFRLKMDP | 1.01 |
| **31** | Training | VCNRIEPLKPIL | 0.95 |
| **32** | Training | GNKKNWRWYKNR | 0.93 |
| **33** | Training | KPWHGCASRTKR | 0.93 |
| **34** | Training | NKAKACNGMKPR | 0.92 |
| **35** | Training | TIKPRMVKLPSV | 1.08 |
| **36** | Training | MKVAAAMAPKQV | 0.94 |
| **37** | Training | NWRDCLSLIVPN | 0.96 |
| **38** | Training | PLNIAISNSPDS | 0.51 |
| **39** | Training | DCLAVPPNAVTI | 0.95 |

| | | | |
|---|---|---|---|
| **40** | Training | VNSDPSNGQMRD | 1.22 |
| **41** | Training | RGKMPLRWMTRK | 0.96 |
| **42** | Training | IKGMNIKMPTDQ | 0.91 |
| **43** | Training | PEGVSWRVLKAN | 0.94 |
| **44** | Training | RPPMLNVVRVVG | 0.92 |
| **45** | Training | YEYEYEYEYEYE | 0.35 |
| **46** | Training | KSKSKSKSKSKS | 0.99 |
| **47** | Training | PKVVKDMPPWDT | 1.24 |
| **48** | Training | MLMKVPVIRRVW | 0.98 |
| **49** | Training | VARVDIDLQAIA | 0.94 |
| **50** | Training | VITNNWALAQVR | 0.96 |
| **51** | Training | TISDIVKVVIRS | 0.96 |
| **52** | Training | KSSKSSKSSKSS | 0.95 |
| **53** | Training | WLLDLRRQMDDT | 1.45 |
| **54** | Training | RDPDKNNRDWPI | 1.06 |
| **55** | Training | LDDDTPDRNRDW | 1.74 |
| **56** | Training | VGQNNDERRRQR | 1.01 |
| **57** | Training | PGPPAGPTLSNR | 1.02 |
| **58** | Training | NSPQLNLNPPQS | 0.99 |
| **59** | Training | NRSMPLPMNPGD | 1.16 |
| **60** | Training | NAARMNRLNDAM | 1.04 |
| **61** | Training | ANNNPWSGMMNG | 0.96 |

| | | | |
|---|---|---|---|
| **62** | Training | MWVKGMKHKKMK | 1.01 |
| **63** | Training | VPKDKRKKLTNP | 0.95 |
| **64** | Training | AACPNLAVAAPM | 0.94 |
| **65** | Training | VPKRLVVVMNTC | 1.06 |
| **66** | Training | EAPMPKVNVIVN | 1.05 |
| **67** | Training | PLPDNVVKAVVW | 1.02 |
| **68** | Training | CKPARTDWPPMP | 1.01 |
| **69** | Training | CATNAKNNNMRG | 1.00 |
| **70** | Training | VINKVISNPCVN | 0.99 |
| **71** | Training | LHSQWLKVDHLL | 1.01 |
| **72** | Training | QTATENSQMNSG | 1.14 |
| **73** | Training | KKKKKKKKKKKK | 1.01 |
| **74** | Training | KSSSKSSSKSSS | 0.96 |
| **75** | POET Motif (Epoch 1) | MDDLTGDNKNWV | 1.44 |
| **76** | POET Motif (Epoch 1) | QDTTMMTRLDTM | 1.06 |
| **77** | POET Motif (Epoch 1) | VDRNTDTDSVDT | 1.98 |
| **78** | POET Motif (Epoch 1) | KVDCDDVDRLDS | 2.00 |
| **79** | POET Motif (Epoch 1) | DMMHDKTDCNDT | 1.71 |
| **80** | POET Motif (Epoch 1) | TCTPTNTLNTMD | 1.23 |
| **81** | POET Motif (Epoch 1) | IDMTCSMTDDDT | 1.00 |
| **82** | POET Motif (Epoch 1) | QTCRNDGTIDKH | 1.10 |
| **83** | POET Motif (Epoch 1) | TNAGTMCTTNTD | 1.17 |

| | | | |
|---|---|---|---|
| **84** | POET Motif (Epoch 1) | CNDDTAGECTND | 2.08 |
| **85** | POET regex (Epoch 1) | CDVDPITCDNDD | 1.95 |
| **86** | POET regex (Epoch 1) | LDNPQEDADDDG | 2.12 |
| **87** | POET regex (Epoch 1) | LEDDIDCDNNDF | 1.77 |
| **88** | POET regex (Epoch 1) | DTMTLNDQETDD | 2.05 |
| **89** | POET regex (Epoch 1) | WHCDQTDDIDGH | 2.04 |
| **90** | POET regex (Epoch 1) | RDDETLDTLDSG | 2.05 |
| **91** | POET regex (Epoch 1) | NSAMADQADDDP | 1.12 |
| **92** | POET regex (Epoch 1) | IDCRGTEDDDPN | 2.30 |
| **93** | POET regex (Epoch 1) | RVDMDDTIDGMH | 1.83 |
| **94** | POET regex (Epoch 1) | NDGGDPMADNDN | 1.92 |
| **95** | EF hand 3 reveresed[2] | EKKDLTGDNDPD | 1.98 |
| **96** | EF hand 4 reveresed[2] | ERADITGDNDPN | 2.08 |
| **97** | Hans EF hand 1[6] | NKDNDDSLEIAE | 1.62 |
| **98** | Hans EF hand 2[6] | NPDGDTTLESGE | 1.64 |
| **99** | Hans EF hand 3[6] | NKDGDQTLEMDE | 1.77 |
| **100** | Hans EF hans 4[6] | DANKDGKLTAAE | 1.14 |
| **101** | POET Motif (Epoch 2) | DEDDDLECVDDA | 2.02 |
| **102** | POET Motif (Epoch 2) | GDLVDCDEDCDD | 1.76 |
| **103** | POET Motif (Epoch 2) | CDEDDEGVKDDD | 2.15 |
| **104** | POET Motif (Epoch 2) | NDGDGPCCEDDD | 2.07 |

| | | | |
|---|---|---|---|
| **105** | POET Motif (Epoch 2) | DDEDDTSGGDTK | 2.11 |
| **106** | POET Motif (Epoch 2) | GDDEDDQCHQDG | 2.17 |
| **107** | POET Motif (Epoch 2) | EDGDVIGSGKDD | 2.07 |
| **108** | POET Motif (Epoch 2) | TEGGTDDTGGND | 2.08 |
| **109** | POET Motif (Epoch 2) | CPTDDGHDDHDD | 2.10 |
| **110** | POET Motif (Epoch 2) | HDDGVGVDGGDG | 1.88 |
| **111** | POET regex (Epoch 2) | CDVDPITCDNDD | 2.14 |
| **112** | POET regex (Epoch 2) | SDDNHSDLGDDL | 2.12 |
| **113** | POET regex (Epoch 2) | FEDHSDDTDEDT | 2.07 |
| **114** | POET regex (Epoch 2) | VDDSDPDDVKDT | 2.10 |
| ***115*** | POET regex (Epoch 2) | DDDSDEDSTDSH | 1.51 |
| ***116*** | POET regex (Epoch 2) | DSDEDTHVDCQE | 2.16 |
| ***117*** | POET regex (Epoch 2) | TQDSDDGMEDED | 2.09 |
| ***118*** | POET regex (Epoch 2) | STDTAGDDATEG | 1.05 |
| ***119*** | POET regex (Epoch 2) | ENHSGQHCHDTG | 1.81 |
| ***120*** | POET regex (Epoch 2) | VCEDHDPNDSHI | 1.88 |

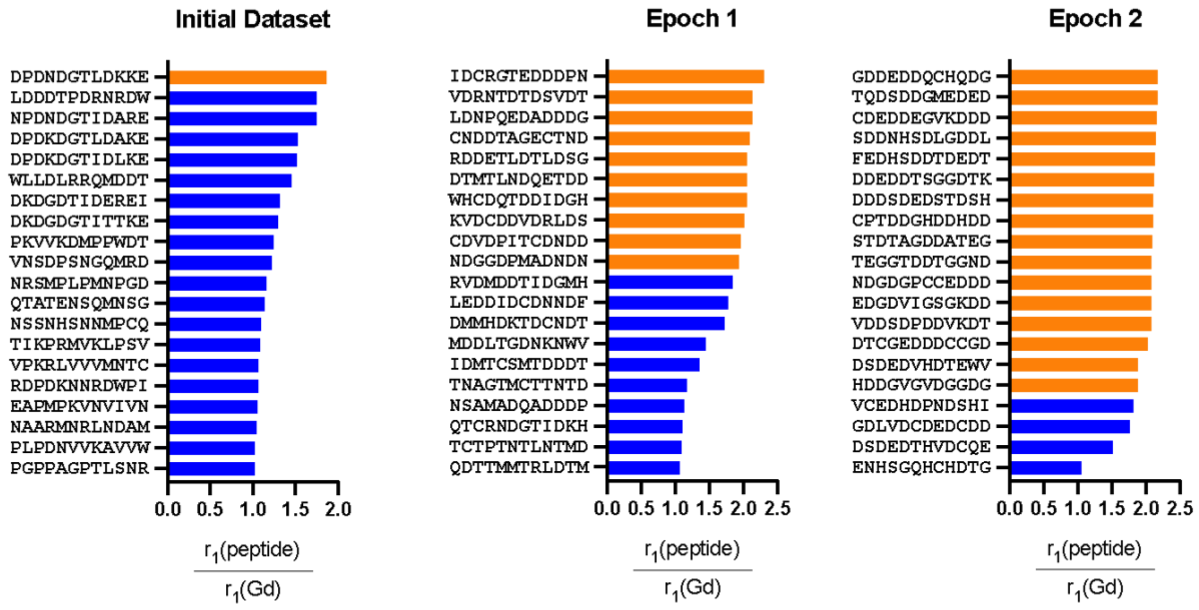


**Figure S2**. The higher the $r_1$ ratio, the better the MRI contrast. $GdCl_3$ is used as an internal control. Left - Partial initial dataset for POET. Middle and right – POET predictions from the initial dataset. Orange bars have an $r_1$ ratio equal to or higher than EF hand 3, the best peptide in the initial dataset.
